# Photocrosslinkable and biodegradable hydrogels for the controlled delivery of exosomes

**DOI:** 10.1101/2025.04.01.646472

**Authors:** Sergio Ayala-Mar, Oju Jeon, Pedro Chacón-Ponce, José González-Valdez, Eben Alsberg

## Abstract

Local and controlled delivery of exosomes using hydrogels can improve the efficacy of exosome-based therapeutics. Hydrogels based on oxidized and methacrylated alginate (OMA) can be produced through photocrosslinking and exhibit controllable degradability. The degradation properties of OMA hydrogels can be tailored by varying the alginate oxidation degree, which is advantageous for exploring new therapeutic strategies. Here, we prepared photocrosslinkable, biodegradable, cytocompatible, and tunable alginate hydrogels for exosome delivery. We synthesized OMA with different degrees of alginate oxidation to generate OMA hydrogels that undergo both rapid and slow degradation. The degradation rate, swelling ratio and mechanical stability of the OMA hydrogels were investigated. Likewise, we examined exosome release rate and kinetics. The cytocompatibility and functionality of these systems were confirmed. Increasing the alginate oxidation degree greatly enhanced the degradation of OMA hydrogels. Rapid degradation correlated with accelerated exosome release, while slow degradation correlated with enhanced exosome retention. Physicochemical properties including the rheological behavior and swelling also influenced the exosome release rate. Kinetic equation fits showed that the Korsmeyer-Peppas model can be used to describe the release process. These findings demonstrate that OMA hydrogels are suitable delivery carriers for the controlled release of exosomes and highlight the potential use of this technology in applications that depend on either rapid or slow exosome delivery and the temporary presence of the hydrogel, including tissue engineering, wound healing and localized and targeted drug delivery.

## 1. Introduction

Extracellular vesicles (EVs) are lipidic particles that naturally transfer bioactive molecules amongst cells and tissues. Exosomes are a subtype of EVs of endosomal origin, which have been recently classified as small EVs (sEVs) because of their size range of ∼10-200 nm. Exosomes are widely proposed for therapeutic applications because of their high stability, low toxicity and drug loading capacity [1]. These applications include tissue microenvironment modulation and delivery of therapeutics [2]. Additionally, enrichment of specific proteins and nucleic acids in exosomes is a promising strategy to target critical signaling pathways related to various diseases, including cancer [3–5].

Specifically, directed drug delivery to overcome therapy resistance and limited response rates, improving antitumor immunity and novel combination therapies are among the clinical applications proposed for exosomes in cancer treatment [6–8]. Nonetheless, the pharmacokinetics and biodistribution of sEVs limit their use in these approaches. Exosomes show a short half-life and high accumulation in the liver, kidneys, spleen, and lungs after intravenous injection [9]. These challenges can be tackled by encapsulating EVs within natural and synthetic hydrogels, which have shown to be effective systems for the controlled release of bioactive molecules at the target site [10,11].

In recent years, hydrogels have become the first choice to develop drug delivery systems because of their cargo loading capacity and biocompatibility [12]. These systems are crosslinked polymer networks that can be used to target specific tissues during an appropriate time. Indeed, injectable hydrogels may enhance the efficacy of therapeutics and prevent side effects through local administration [10,13].

Different hydrogels offer varied sEV release profiles depending on their nature and properties. Likewise, exosomes can be integrated into hydrogels through multiple approaches, either by being added to preformed hydrogels or during the gelation process [14,15]. The choice of method depends on several factors, including the application intended for the hydrogel system, the properties of the polymer, and the specific gelation mechanism. Currently, the main method involves using polymer solutions prepared with an appropriate viscosity, suitable for incorporating exosomes before gelation [16].

Thermogelation is one the most widely used strategies in healthcare to produce gels, primarily using poloxamers, such as Pluronic F127, that form hydrogels at body temperature. In fact, Pluronic F127 has been combined with F68, to successfully incorporate platelet-rich plasma-derived EVs for continuous release over 28 days, demonstrating therapeutic potential in delaying the development of osteoarthritis in mice [17]. Similarly, polycitrate-PEI combined with F127 formed a hydrogel that provided long-term retention and sustained release of EVs over 50 days [18].

Thermogelation can also be achieved using natural polymers like chitosan, collagen, and cellulose, combined with salts. Chitosan hydrogels containing exosomes, for instance, have been used for corneal epithelium repair in rat models [19]. Collagen-based patches containing iPSCs-derived exosomes have shown significant reduction in infarct size in rats, while methylcellulose-based hydrogels have proven effective in treating critical limb ischemia by delivering EVs derived from endothelial cells [20,21].

On its part, photocrosslinking offers another approach for generating hydrogels loaded with exosomes. For example, methacrylated gelatin (GelMA) based hydrogels have successfully incorporated HUVEC-derived EVs, during photocrosslinking, showing sustainable release over 7 days *in vitro* [22]. When applied cutaneously, the GelMA hydrogel loaded with exosomes accelerated re-epithelialization, promoted collagen maturity, and improved angiogenesis, in a wounded rat model.

In addition to thermoresponsive and photocrosslinked hydrogels, hydrogels based on physically crosslinked polymers also demonstrate considerable potential in exosome delivery. For example, hydrogels can be formed through Schiff base linkages by mixing aldehyde-modified methylcellulose and chitosan grafted PEG forms. This particular formulation was used for the controlled delivery of placental MSC-derived EVs in a diabetic rat model, achieving full skin structure and function recovery within 15 days [23].

Together, these diverse hydrogel systems demonstrate the potential of using natural and synthetic polymers for effective exosome delivery in various therapeutic contexts, but other materials present advantages that can be harnessed in sEV delivery applications. A good example is alginate, a natural polymer that has been widely used as an effective biomaterial for bioactive delivery because of its safety and stability [24]. Previous studies have shown that exosomes can be encapsulated in alginate-based hydrogels for controlled release, while preserving their properties and function. [25,26].

Unmodified alginate hydrogels, typically formed through ionic crosslinking, exhibit a range of physical properties based on their formulation and preparation conditions. These properties can be tuned, including optimization of the polymer concentration and molecular weight, to meet specific application requirements. Furthermore, the degradation rate of alginate hydrogels can be effectively controlled by chemical modifications, such as oxidation with sodium periodate [27]. This approach allows varying the alginate oxidation degree to control its degradability [28]. Degradability is an important factor for bioactive delivery and biodegradable hydrogels do not need to be removed after the exosomes have been released.

Alginate can also be modified by methacrylation, which allows gelation after light exposure. Besides the ease of this crosslinking process, photocrosslinked alginate hydrogels exhibit substantial advantages over ionically crosslinked hydrogels, including precise spatial and temporal control over the gelation process and mild crosslinking conditions [29,30]. For instance, photocrosslinkable alginate hydrogels have shown tailorable physicochemical properties and can be delivered in a minimally invasive manner [29]. Indeed, the mechanical properties of oxidized and methacrylated alginate (OMA) hydrogels can be tuned by varying the alginate oxidation and methacrylation degrees, macromer concentration, and crosslinking density [31].

As a result, OMA hydrogels represent a significant advancement in the field, offering enhanced control over degradation rate, safety, and tunability. These properties make OMA hydrogels promising candidates for a wide range of therapeutic applications, effectively addressing key limitations observed in other hydrogel-based sEV delivery systems [16,32]. For instance, thermogels face challenges related to precise control of mechanical properties and degradation rates. Similarly, photocrosslinkable hydrogels, such as GelMA-based hydrogels can face limitations in terms of degradability.

In contrast, the degradation rate of OMA hydrogels is finely tunable, which might allow a precise control over exosome release kinetics, a significant improvement over systems where degradation rates are less predictable [33–35]. Even more, the mechanical properties of OMA hydrogels are also highly adaptable and therefore these can be customized to suit specific tissue targets and delivery needs, offering a level of versatility that is not as readily achievable in many other systems [36].

Based on these observations, during this work, photocrosslinkable OMA hydrogels were produced with different alginate oxidation degrees. It was anticipated that the alginate oxidation degree could be used to tune the degradation rate of the gels. Accordingly, the degradation rate, swelling kinetics and mechanical stability of the OMA hydrogels was assessed. Then, cancer cell-derived exosomes were encapsulated in OMA hydrogels designed to undergo rapid or slow degradation. Subsequently, the exosome release rate and kinetics were assessed. In addition, we evaluated the cytocompatibility of these systems and their ability to deliver functional exosomes *in vitro*.

Our results show that rapid degradation of the hydrogels results in accelerated exosome release, while slow degradation increases exosome retention within the hydrogels. In this context, alginate-based hydrogels might be used as a controlled release depot, prolonging the release of exosomes at the target site, or as a delivery vehicle for rapid exosome release, which can modulate acute biological processes during a short period [11,37]. By addressing the current limitations in sEV delivery, OMA hydrogels could significantly advance therapeutic strategies in areas such as cancer treatment, tissue regeneration, and targeted drug delivery.

## 2. Materials and methods

### 2.1 Synthesis of oxidized and methacrylated alginate (OMA)

OMA with a theoretical 2% (2-OMA) and 5% (5-OMA) oxidation degree and a theoretical 20% methacrylation degree were synthesized according to reported literature protocols [29,36]. The feeding ratios of reactants and reagents to synthetize OMA are shown in **Table 1**. Briefly, sodium alginate [10 g, Protanal LF120 M LVG low viscosity (157 mPa·S at 1 % water, FMC Biopolymer)] was dissolved in 900 mL of ultrapure deionized water (diH_2_O) overnight. To synthesize oxidized alginate (OA), sodium periodate (0.218 or 0.545 g, Sigma) was dissolved in 100 mL diH_2_O, added to the alginate solution under stirring in the dark at room temperature (RT) and then allowed to react for 24 h.

**Table 1.** Reactant and reagent feeding ratios for the synthesis of OA and OMA.

| Polymer | Alginate (g) | NaIO<sub>4</sub> (g) | MES/NaCl (g/g) | NHS/EDC (g/g) | AEMA (g) |
| --- | --- | --- | --- | --- | --- |
| 2-OMA | 10 | 0.218 | 19.52/17.53 | 1.118/3.888 | 1.688 |
| 5-OMA | 10 | 0.545 | 19.52/17.53 | 1.118/3.888 | 1.688 |

To synthesize OMA, 2-morpholinoethanesulfonic acid (MES, 19.52 g, Sigma) and NaCl (17.53 g, Sigma) were directly added to an OA solution (1 L), and pH was adjusted to 6.5 with 5 N NaOH (Sigma). Then, *N*-hydroxysuccinimide (NHS, 1.118 g; Sigma) and 1-ethyl-3-(3-dimethylaminopropyl)-carbodiimide hydrochloride (EDC, 3.888 g; Sigma) (molar ratio of NHS:EDC = 1:2) were added to the mixture. After 10 min, 2-aminoethyl methacrylate (AEMA, 1.688 g, Polysciences) (molar ratio of NHS:EDC:AEMA = 1:2:1) was added to the solution, and the reaction was maintained in the dark at room temperature (RT) for 24 h. Then, acetone (OMA 1: acetone 3) was added to precipitate the reaction mixture for 10 min. The precipitate was dried in a fume hood and rehydrated to a 1% (w/v) solution in diH_2_O for further purification. OMA was purified by dialysis against diH_2_O (MWCO 3500, Spectrum Laboratories Inc.) for 3 days at 4 °C, treated with activated charcoal (5 g/L, 100 mesh, Oakwood Chemical) for 30 min, filtered (0.22 μm filter) and lyophilized. OMA was stored at -80 °C until further use.

To generate photocrosslinked OMA hydrogels, 2-OMA (2% and 4% w/v) and 5-OMA (4% and 8% w/v) were dissolved in Dulbecco’s Modified Eagle Medium (DMEM (Sigma) with a photoinitiator [2-Hydroxy-4’-(2-hydroxyethoxy)-2-methylpropiophenone, 0.05% w/v, Sigma] at pH 7.4. OMA solutions were placed between two quartz plates separated by 1 mm spacers and photocrosslinked with UV light (OmniCure Series 1000, EXFO, Canada) at 20 mW/cm^2^ for 60 s to form hydrogels.

### 2.1 Characterization of swelling kinetics, degradation rates and mechanical properties of OMA hydrogels

To determine the swelling ratio of the hydrogels, photocrosslinked OMA hydrogels (with an approximate diameter of 3.5 cm) were prepared as described above. Then, the samples were lyophilized, and dry weights (W_i_) were measured. Dried hydrogel samples were immersed in 15 mL of DMEM with a photoinitiator at pH 7.4 and incubated at 37 °C to reach equilibrium. At predetermined time points, the samples were removed, and the swollen (W_s_) hydrogel weights were measured. The swelling ratio (Q) was calculated as follows: Q = W_s_/W_d_.

To determine the degradation rates, the dried hydrogel samples were weighed (W_i_) and incubated in 15 mL of DMEM with photoinitiator at pH 7.4 at 37 °C. At predetermined time points, the medium was removed, and the hydrogel samples were collected, lyophilized, and weighed (W_d_). The percent mass loss was calculated as follows: Percentage mass loss = (W_i_ − W_d_)/W_i_ × 100.

Dynamic rheological examination of the OMA hydrogels was performed to evaluate their mechanical properties with a Kinexus ultra + rheometer (Malvern Panalytical, UK). In oscillatory mode, a parallel plate (8 mm diameter) geometry measuring system was employed, and the gap was set to 0.9 mm. Briefly, OMA solutions were placed between two quartz plates with 1 mm spacers and photocrosslinked as described above. Then, hydrogel disks were created using a biopsy punch (d = 8 mm). Photocrosslinked OMA hydrogel disks were placed between the plates, all the tests were started at 25 ± 0.1 °C, and the plate temperature was maintained at 25°C. Oscillatory frequency sweep (0.1–10 Hz at 1 % shear strain) tests were performed to measure storage (G′) and loss (G″) moduli.

### 2.2 Cell culture

Human cervical adenocarcinoma (HeLa) cells were obtained from the American Type Culture Collection (ATCC, USA). HeLa were cultured in DMEM supplemented with 10% fetal bovine serum, 100 U/mL penicillin and 100 µg/mL streptomycin and incubated at 37 °C in a humidified 5% CO_2_ incubator.

HeLa spheroids were generated in a 96-well round bottom Ultra Low Attachment plate (Corning, USA). Cells were seeded at a density of 1000 cells/well, after which plates were centrifuged at 250 *x g* for 5 min and incubated for 4 days prior to treatment. Cell culture medium was refreshed every other day. To isolate exosomes, 5×10^6^ cells/flask were seeded in T175 flasks and incubated for 24 h in complete growth medium. Then, cells were washed 3 times with PBS (Sigma-Aldrich, MO) and incubated for 48 h in serum free medium under standard culture conditions.

### 2.3 Exosome isolation

Conditioned medium was harvested, and the ExoQuick-TC reagent (System Biosciences, USA) was used to isolate exosomes according to the manufacturer’s protocol with slight modifications [38]. Briefly, 30 mL of conditioned medium was first centrifuged at 3,000 *x g* at 4 °C for 15 min. The supernatant was collected and centrifuged at 10,000 *x g* at 4 °C for 30 min. The supernatant was subsequently filtered (0.22 μm filter) and concentrated using 100-kDa molecular weight cutoff Amicon Ultra-15 centrifugal filter units (Merck, Germany) at 4,000 *x g* at 4 °C for 45 min. Then, 10 mL of concentrated conditioned medium were mixed with 2 mL of the ExoQuick-TC reagent by vortexing. The sample was incubated overnight at 4 °C and centrifuged at 1,500 *x g* for 30 min to pellet the exosomes. The supernatant was discarded, and the pellet was resuspended in PBS.

Protein concentration of the exosome samples was determined with the Micro BCA protein assay kit (Pierce, Thermo Fisher Scientific, USA) according to the manufacturer’s instructions. A schematic representation of this process and the initial exosome identification results of the obtained HeLa-derived sEVs can be found on **Fig. S1A** in Supplementary Information.

### 2.4 Characterization of exosome particle concentration, size, morphology, and biochemical markers

To determine exosome particle concentration, size distribution and mean size, nanoparticle tracking analysis (NTA) was performed by using a NanoSight NS300 (Malvern Instruments, UK) instrument fitted with a sCMOS camera and a blue laser (488 nm). Briefly, sEV samples were diluted in PBS (1:1000) to an appropriate concentration within the linear range of the instrument, vortexed and filtered (0.22 µm). For each sample, at least 5 videos of 30 s were recorded to generate enough completed tracks (>200) for an accurate size distribution. Capture settings were set as follows: Screen gain: 1; Camera level: 15; Slider shutter: 1206; Slider gain: 366; FPS: 25; Syringe pump speed: 100. Video processing settings for analysis were set as follows: Detection threshold: 5; Blur size: Auto; Max jump distance: Auto. Data was analyzed using the NTA 3.2 software.

The morphology of the exosomes was observed by transmission electron microscopy (TEM). Briefly, 10 μL of an exosome solution sample were loaded onto a formvar-coated copper grid. Then, the exosomes were negatively stained with 2% uranyl acetate for 5 min. After being air-dried, the grids were visualized by TEM (JEOL JEM-ARM200CF, JOEL, Japan) at 80 kV.

For the detection of CD63 (a widely used and recognized exosomal biomarker) on isolated exosomes, an ELISA for CD63 (System Biosciences, USA, Cat. No. EXEL-ULTRA-CD63-1) detection kit was used according to the manufacturer’s recommendations. Acetylcholinesterase (AChE) activity was measured in lysed exosome samples using an AChE activity assay kit (System Biosciences, USA, Cat. No. FCET96A-1), following the manufacturer’s protocol.

### 2.5 Exosome labelling and encapsulation in OMA hydrogels

Fluorescent labeling of exosomes was carried out using the BODIPY TR ceramide dye (Thermo Fisher Scientific, USA) according to manufacturer’s instructions [39]. Briefly, 100 µL of an exosome sample corresponding to 100 µg of protein were mixed with 1 µL of a 1 mM solution of BODIPY TR ceramide by vortexing and incubating the mixture for 1 h at 37 °C. Excessive dye from labeled exosomes was removed using a Vivaspin 500 centrifugal concentrator (Thermo Fisher Scientific, USA) with a 3,000 molecular weight cutoff filter. Labeled exosomes were immediately used after staining.

To encapsulate exosomes in OMA hydrogels, exosomes suspended in PBS were added to the hydrogel solution (1:20 volume ratio) and mixed by gentle pipetting before crosslinking. Then, OMA solutions were photocrosslinked with UV light to form hydrogels, as described above. Exosomes were encapsulated within OMA hydrogels, achieving an exosomal protein concentration of 50 µg per mL of OMA hydrogel solution, equivalent to approximately 5×10^9^ particles/mL.

### 2.6 Exosome release and kinetics from OMA hydrogels

To quantify sEV release from OMA hydrogels, an *in vitro* release assay was performed. For this purpose, BODIPY-TR ceramide labeled exosomes were added to hydrogel solutions. Then, 30 µL of each sample were crosslinked as described above and placed in a 1.5 mL conical tube (Eppendorf, Germany). 300 µL of release medium consisting of DMEM (Sigma) with a photoinitiator at pH 7.4 were added to each tube and incubated at 37 °C in 5% CO_2_. At predetermined time intervals, the release medium was replaced with fresh solution to maintain the total volume.

To quantify the number of exosomes in the medium, 300 µL of release medium were placed in each well of an opaque 96-well (Corning, USA) plate and fluorescence intensity was measured (Excitation/Emission: 589 nm/615nm) using a microplate reader (BioTek, USA). The content of the released exosomes was determined from the fluorescence intensity of the solution according to a standard curve.

The exosome released percentage was determined by the following equation:

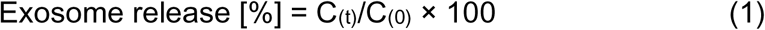

where C_(0)_ and C_(t)_ represent the amount of exosomes loaded and the amount of exosomes released at time (t), respectively.

To study the exosome release process from the OMA hydrogels, the release data was fitted to various kinetic models to describe the mechanism of release. This analysis included zero-order kinetics [40,41], first-order kinetics [42], the Higuchi’s square root of time equation or diffusion model [43], and the Power law equation, also known as the Korsmeyer-Peppas model or diffusion/relaxation model [44].

Accordingly, the exosome release data obtained as described above were correlated according to the following kinetic equations:

Zero-order kinetics

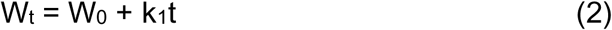

First-order kinetics

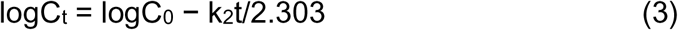

Higuchi’s square root of time equation (diffusion model)

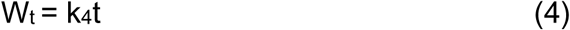

Power law equation (diffusion/relaxation model)

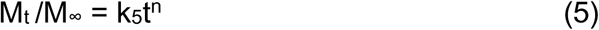

where k is the exosome release constant; C_t_ and C_0_ are the exosome concentrations at the initial time and testing time; W_t_ and W_0_ represent the exosomal mass at the initial time and testing time, as reflected by the cumulative release percentage; M_t_/M_∞_ is the fractional exosome release into the release medium; and n is the diffusional constant that characterizes the exosome release transport mechanism. Data from five independent replicates were used to estimate the average correlation coefficient to each of the previously described kinetic models by using Excel Solver Software (Microsoft, USA).

### 2.7 Biocompatibility and functionality of OMA hydrogels

To evaluate the viability of HeLa cells during coculture with OMA hydrogels, we performed a Transwell assay using 24-well Transwell chambers with 8 μm polycarbonate membranes (Millipore, MA, USA). HeLa cells (1×10^5^ cells/well) were seeded in the lower chamber of a Transwell (Corning, USA) tissue culture plate and incubated for 24 h under standard culture conditions. Then, 100 µL of photocrosslinked 2-OMA or 5-OMA (4% w/v) hydrogels were placed in the upper chamber insert of the Transwell tissue culture plate and incubated at 37 °C in a humidified 5% CO_2_ incubator.

At predetermined time points, the insert was removed and Live/Dead staining consisting of fluorescein diacetate (FDA, Sigma) and ethidium bromide (EB, Fisher Scientific) was used to assess viability. Shortly, 20 μL of staining solution, prepared by mixing 1 mL FDA solution (1.5 mg/mL in dimethyl sulfoxide), 0.5 mL EB solution (1 mg/mL in PBS) and 0.3 mL PBS, were added into the lower chamber and incubated for 15 min at RT. Then, HeLa cells were imaged using a fluorescence microscope (ECLIPSE TE 300) equipped with a digital camera (MU1402-NI05, AmScope). Cell viability was calculated as follows: [number of green (live) stained cells]/ (number of green (Live) + red (Dead) stained cells) X 100%.

To assess exosome delivery from OMA hydrogels, HeLa (1×10^5^ cells/well) were seeded in the lower chamber of a 24-well Transwell tissue culture plate, as described above. Then, 100 µL of photocrossslinked 2-OMA or 5-OMA (4% w/v) hydrogels containing fluorescently labeled exosomes were placed in the upper chamber insert of the Transwell tissue culture plate, and then the plate was placed in the incubator.

At predetermined time points, the inserts containing the OMA hydrogels were removed, and cells in the lower chamber were fixed with 4% paraformaldehyde solution (Sigma, USA) for 5 min in the dark at RT and washed with PBS twice. Then, cells were imaged as described above to obtain representative images. Fluorescence intensity was measured (Excitation/Emission: 589/615 nm) using the microplate reader. Control samples were used to subtract the average background signal from the experimental groups.

To investigate the viability of HeLa in OMA hydrogels, HeLa spheroids were generated using ULA round bottom plates as described above and encapsulated in OMA hydrogels. Briefly, cell culture medium was completely removed and replaced by 100 µL of either 2-OMA or 5-OMA (4% w/v) hydrogel solutions. Then, the OMA solutions containing the spheroids were photocrosslinked by placing the 96-well plate under UV light at 20 mW/cm^2^ for 60 s. 200 µL of fresh medium were then added to each well, and plates were cultured in the incubator for 3 days. After 3 days, 4 µL of Live/Dead staining solution were added to each well, plates were incubated for 20 min in the dark at RT, and each well was imaged as described above.

To assess exosome delivery to HeLa cells in OMA hydrogels, HeLa spheroids were encapsulated in 2-OMA or 5-OMA (4% w/v) hydrogel solutions containing BODIPY-TR ceramide labeled exosomes and photocrosslinked as described above. 200 µL of fresh medium were added to each well, and plates were cultured in the incubator for 3 days. After 3 days, medium was removed and 200 µL of PBS were added to each well, and then each well was again imaged as described above.

### 2.8 Statistical analysis

Quantitative data was expressed as mean ± standard deviation (SD). The analyzed sample size for data quantification is indicated in the corresponding figure legends. Statistical analysis between two groups was performed using unpaired two-tailed Student’s *t* test. Statistics analysis among multiple groups were compared using a one-way analysis of variance (ANOVA) with the Tukey significant difference post hoc test. GraphPad Prism 6.0 (Inc., USA) was used for statistical analyses and graphics generation. A value of *p* < 0.05 was considered statistically significant.

## 3. Results

### 3.1 Generation of photocrosslinkable and biodegradable OMA hydrogels for exosome delivery

To generate photocrosslinkable and biodegradable hydrogels for exosome delivery, we encapsulated HeLa-derived exosomes in OMA hydrogels with different degrees of alginate oxidation. A general schematic of the OMA synthesis and the process to encapsulate exosomes in OMA hydrogels is shown in **Fig. 1A** and **1B**, respectively. **Fig. 1C** shows 2-OMA and 5-OMA samples before and after crosslinking. Before crosslinking, the samples were in liquid state and appeared fluid. After crosslinking, the samples were solid, regardless of the degree of alginate oxidation.

**Figure 1.**
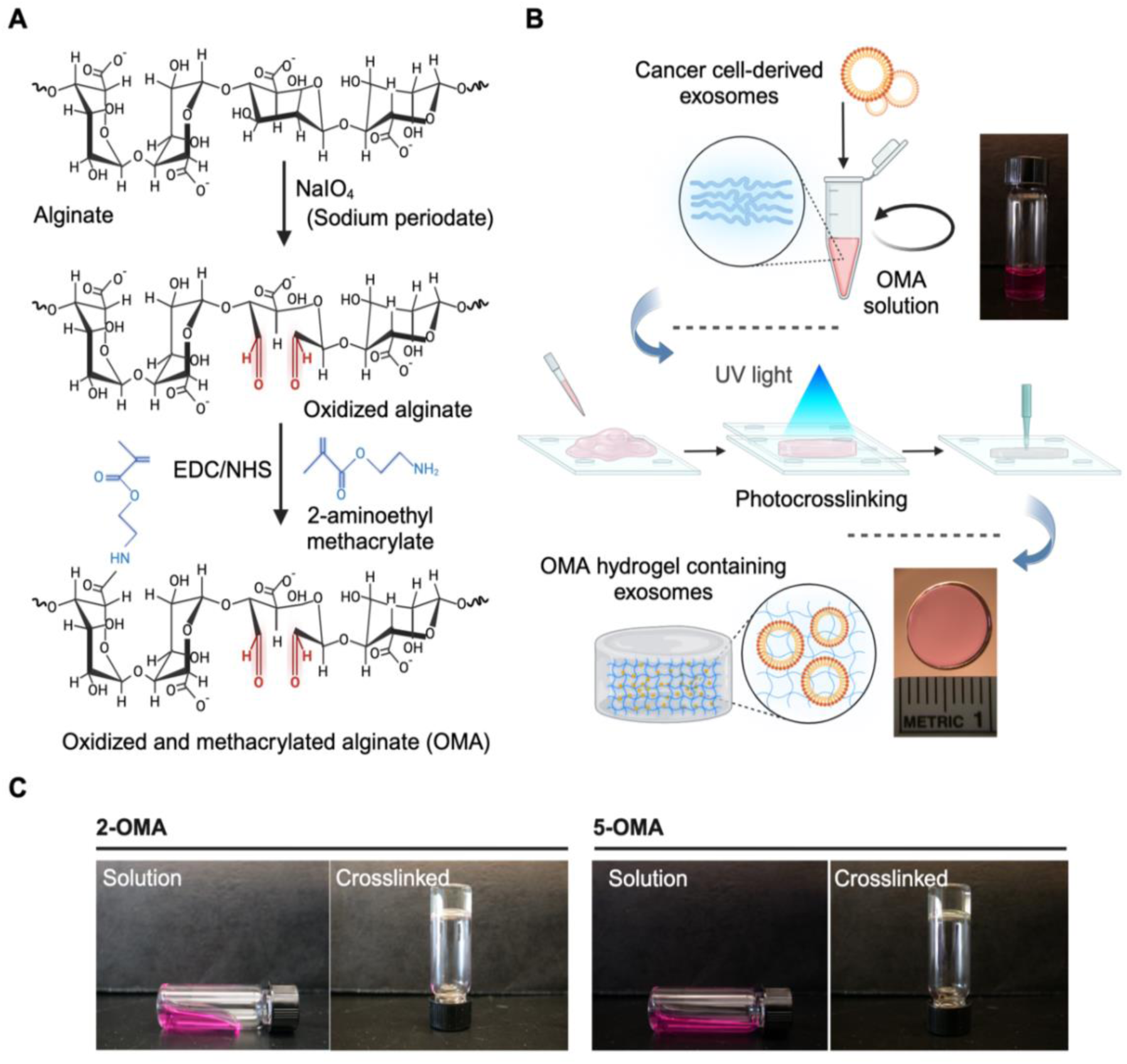
Schematics of OMA hydrogel synthesis for exosome encapsulation. **(A)** Chemical pathway illustrating the sequential modification of alginate through oxidation, followed by methacrylation, resulting in the synthesis of OMA. **(B)** Schematic representation of the encapsulation of exosomes within OMA hydrogels. Initially, exosomes are mixed with the OMA solution. The mixture is then placed between two quartz plates and exposed to UV light to initiate the photocrosslinking process, leading to the formation of a hydrogel matrix that stably encapsulates nanovesicles. **(C)** Comparative optical images show the OMA hydrogels before and after photocrosslinking. The left images show the hydrogel in its initial liquid state, while the right images demonstrate gelation after crosslinking.

To assess the physicochemical properties of these photocrosslinked OMA hydrogels, the degradation rates and the swelling ratios of 4% (w/v) 2-OMA and 5-OMA samples were measured over time. 5-OMA hydrogels exhibited a fast degradation rate with a mass loss of ∼100% by day 6, while the 2-OMA hydrogels showed a much slower degradation rate with a mass loss of only ∼20% after 1 week (**Fig. 2A**). Likewise, the 5-OMA hydrogels showed much faster swelling kinetics compared to 2-OMA hydrogels overtime **(Fig. 2B)**. The swelling ratio of 5-OMA rapidly increased within 5 days and then rapidly decreased. Conversely, the swelling ratio of 2-OMA hydrogels showed a rapid increase during the first day and then a steady increment over the course of 1 week. These results show that 5-OMA hydrogels exhibited rapid degradation and an greater swelling, while 2-OMA hydrogels were slowly degrading with steady increments in swelling over prolonged periods. Indeed, the higher swelling ratio and the faster swelling kinetics observed in the 5-OMA hydrogels are in part due to its faster degradation rate.

**Figure 2.**
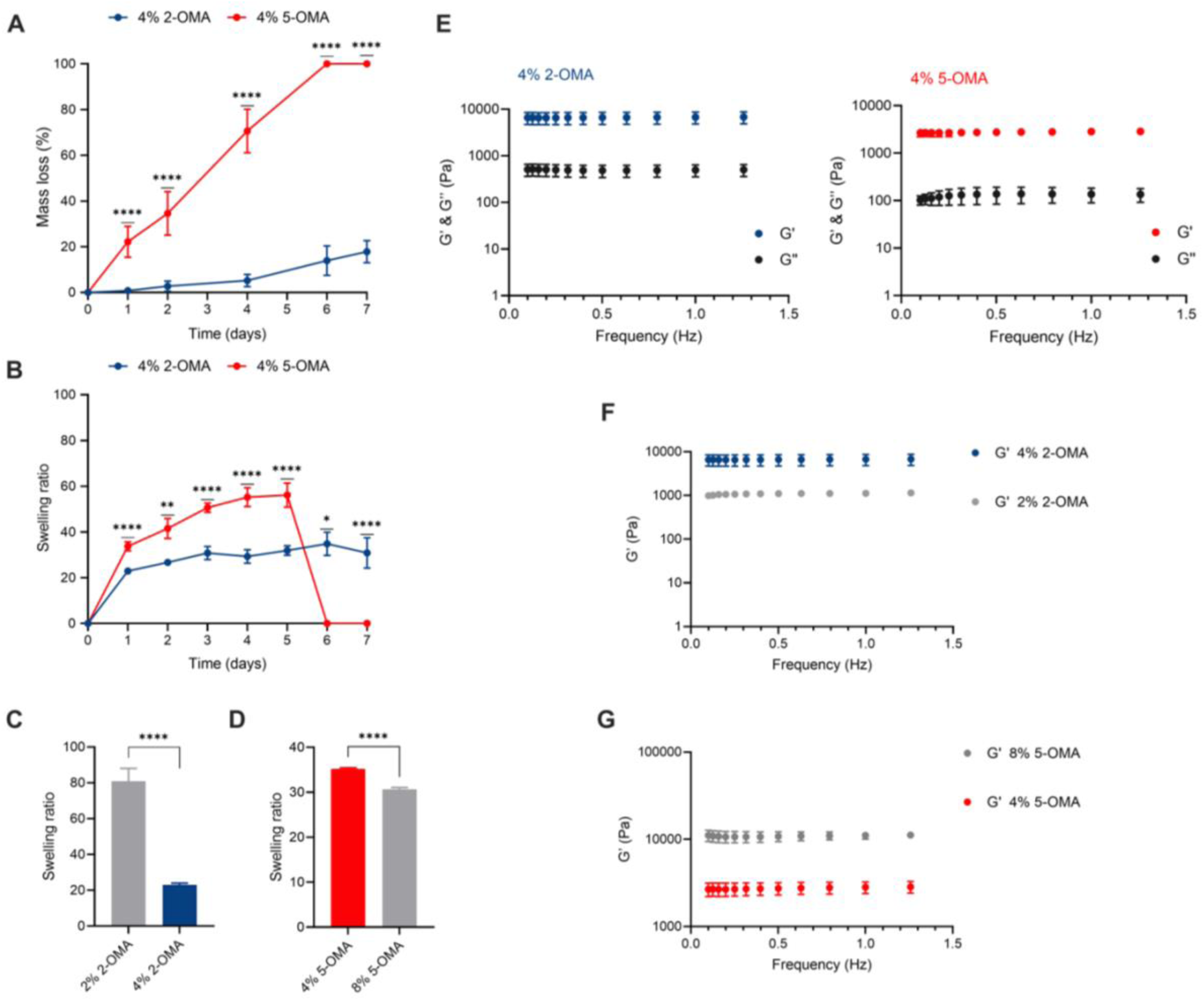
Characterization of OMA hydrogels: degradability, swelling and rheological properties. **(A)** Degradation behavior and **(B)** swelling ratio of OMA hydrogels over time (n=4). Swelling ratio of **(C)** 2-OMA and **(D)** 5-OMA hydrogels having different macromer concentrations after 24 h (n=4). (E) Storage and loss moduli (G’ and G’’) of 2-OMA and 5-OMA hydrogels, and storage modulus of 2-OMA **(F)** and 5-OMA **(G)** hydrogels having different macromer concentrations (n=3). Graphs show the mean ± SD i. Statistical significance between two groups was determined using an unpaired two-tailed Student’s t test (\**p*<0.05, \*\**p*<0.005; \*\*\**p*<0.0005; ****p<0.0001).

To measure the effect of macromer concentration on the swelling ratio of the OMA hydrogels, 2% and 4% (w/v) 2-OMA, and 4% and 8% (w/v) 5-OMA photocrosslinked hydrogels were prepared. After 24 h, the swelling ratio of 2% (w/v) 2-OMA hydrogels was higher when compared to 4% (w/v) 2-OMA hydrogels (**Fig. 2C**). Similarly, the swelling of the 4% (w/v) 5-OMA hydrogels was higher when compared to the 8% (w/v) 5-OMA hydrogels (**Fig. 2D**). Therefore, the swelling of OMA hydrogels appeared to be inversely related to macromer concentration.

Frequency sweep tests were performed to measure the storage (G’) and loss (G’’) moduli of 2-OMA and 5-OMA hydrogels (**Fig. 2E**). G’ was larger than G’’ over the measured frequency range and both moduli exhibited frequency independence in 4% (w/v) photocrosslinked 2-OMA and 5-OMA hydrogels, indicating that they were mechanically stable. G’ and G’’ were larger in 2-OMA hydrogels when compared to the 5-OMA hydrogels at the same macromer concentration, demonstrating that a lower alginate oxidation degree resulted in stiffer hydrogels. These findings suggest that changes in the rheological properties of the hydrogels could be associated with their swelling and degradability under these conditions. To evaluate the macromer concentration effect on the rheological properties of OMA hydrogels, frequency sweep tests of photocrosslinked 2% (w/v) 2-OMA and 8% (w/v) 5-OMA hydrogels were also conducted. G’ and G’’ decreased in the 2% (w/v) 2-OMA hydrogels when compared to 4% (w/v) hydrogels (**Fig.2F**). Similarly, G’ and G’’ were decreased in the 4% (w/v) 5-OMA hydrogels when compared to the 8% (w/v) hydrogels (**Fig. 2G**). Thus, increasing macromer concentration increased the stiffness of the hydrogels, regardless of the alginate oxidation degree. Overall, G’ and G’’ increased with macromer concentration and decreased with alginate oxidation degree.

### 3.2 Quantification and kinetics of exosome release from OMA hydrogels

Exosomes isolated from HeLa cell conditioned medium showed a mean particle size of 121.87 ± 6.09 nm and a mode size of 101.70 ± 10.25 nm. The sample size distribution covered the expected size range of sEV subtypes (**Fig. 3A**) [45,46]. Exosomes were maintained as a stock at a total protein concentration of 1.0 ± 0.016 µg/µL and a particle concentration of 1×10^11^± 7.8×10^10^ particles/mL determined by a MicroBCA protein assay and nanoparticle tracking analysis (NTA), respectively. Particle morphology was assessed by transmission electron microscopy (TEM), which revealed cup-shaped vesicles within the expected size range (i.e., 40-160 nm diameter) (**Fig. 3B**) [45,46]. Additionally, ELISA results confirmed the presence of tetraspanin CD63, and an AChE activity assay indicated exosomal enzymatic activity (**Fig. S1B** and **S1C**, Supplementary Information).

**Figure 3.**
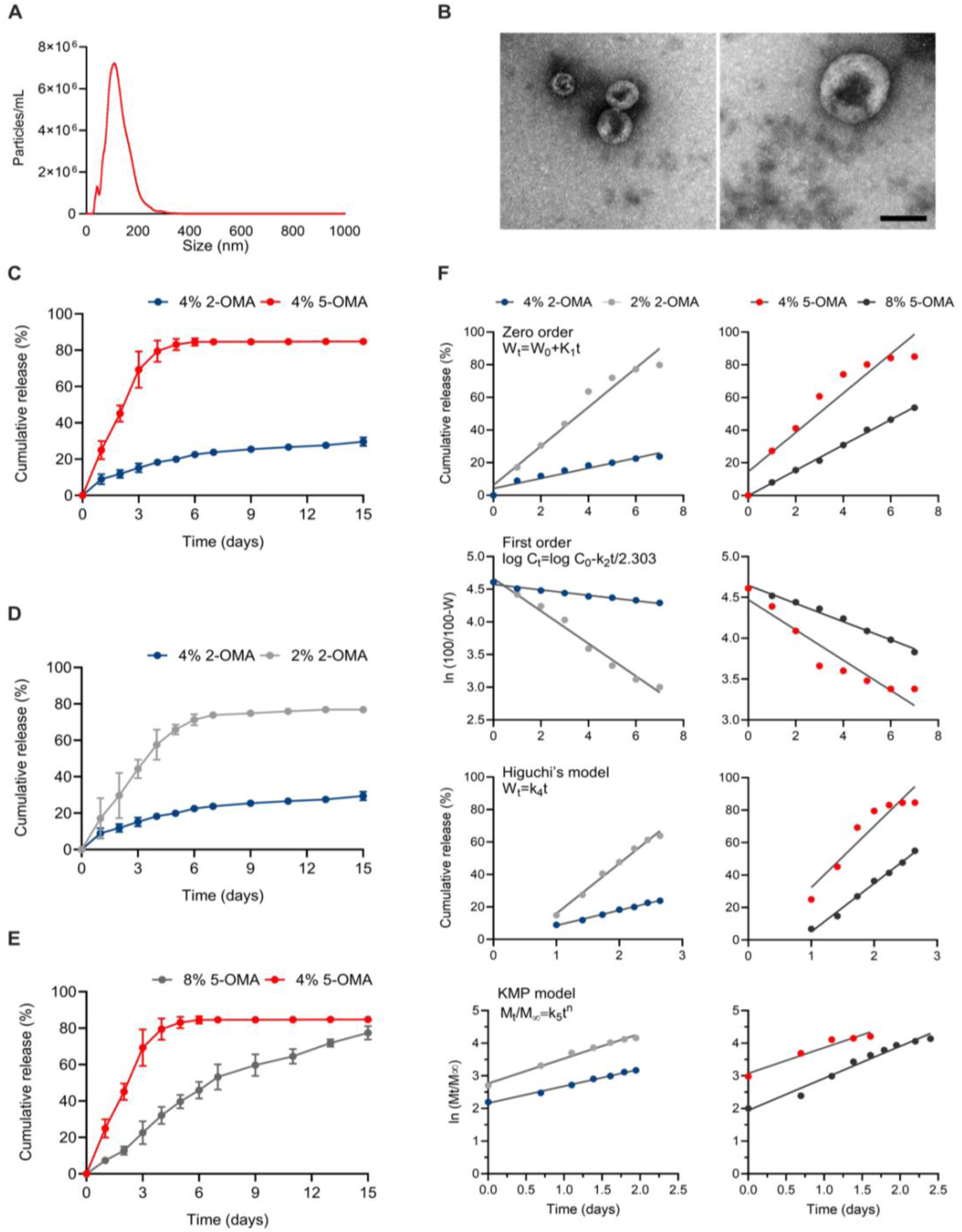
Quantification of exosome release and kinetic modeling. **(A)** Representative particle size distribution of HeLa-derived exosomes by NTA and **(B)** TEM images depicting vesicle morphology (magnification: 50,000x, scale bar: 100 nm). **(C)** Cumulative release profiles of exosomes from 2-OMA and 5-OMA hydrogels. Cumulative release profiles of exosomes from **(D)** 2-OMA and **(E)** 5-OMA hydrogels having different macromer concentrations, and **(F)** kinetic equations fits to different models based on the exosome release data from (2% and 4% w/v) 2-OMA and (4% and 8% w/v) 5-OMA hydrogels. Data indicates the mean ± SD in C-E.

Considering the importance of swelling, degradation rates and mechanical properties in determining the release dynamics of bioactive molecules from hydrogels, we encapsulated exosomes in 4% (w/v) photocrosslinked 2-OMA and 5-OMA hydrogels. This was done to assess how variations in the alginate oxidation degree influence the release profile of these sEVs (**Fig. 3C**). The 2-OMA hydrogels showed a slower release profile compared to the 5-OMA groups at the same macromer concentration. Approximately 15% of the exosomes were released from the 2-OMA hydrogels after 1 week, whereas ∼70% of the exosomes were released from the 5-OMA hydrogels after only 3 days. It is shown that increasing the oxidation degree of alginate hydrogels increases the exosome release rate. Additionally, exosomes released from both types of hydrogels showed a consistent particle size over a 7-day period, indicating that the exosomes retain their size during the dynamic release process (**Fig. S2**, Supplementary Information).

To determine the effect of macromer concentration on the exosome release rate from OMA hydrogels, fluorescently labeled exosomes were encapsulated in 2% and 4% (w/v) photocrosslinked 2-OMA hydrogels (**Fig. 3D**) and 4% and 8% (w/v) photocrosslinked 5-OMA hydrogels (**Fig. 3E**). After 1 week, approximately 80% of the exosomes had been released from 2% (w/v) 2-OMA hydrogels, similar to the 8% (w/v) 5-OMA hydrogels from which around 80% of the exosomes were released after 2 weeks. These results showed the ability to tune the exosome release rate of OMA hydrogels by varying macromer concentration. Accordingly, increasing macromer concentration enhanced exosome retention, while decreasing it accelerated exosome release.

Furthermore, to characterize the exosome release mechanism from OMA hydrogels, equation fits to different kinetic models were calculated from the release data of 2 – 4% (w/v) 2-OMA and 4 – 8% (w/v) 5-OMA photocrosslinked hydrogels (**Fig. 3F**). The regression coefficients of the release process according to the kinetic models was also determined (**Table 2)**.

**Table 2.**
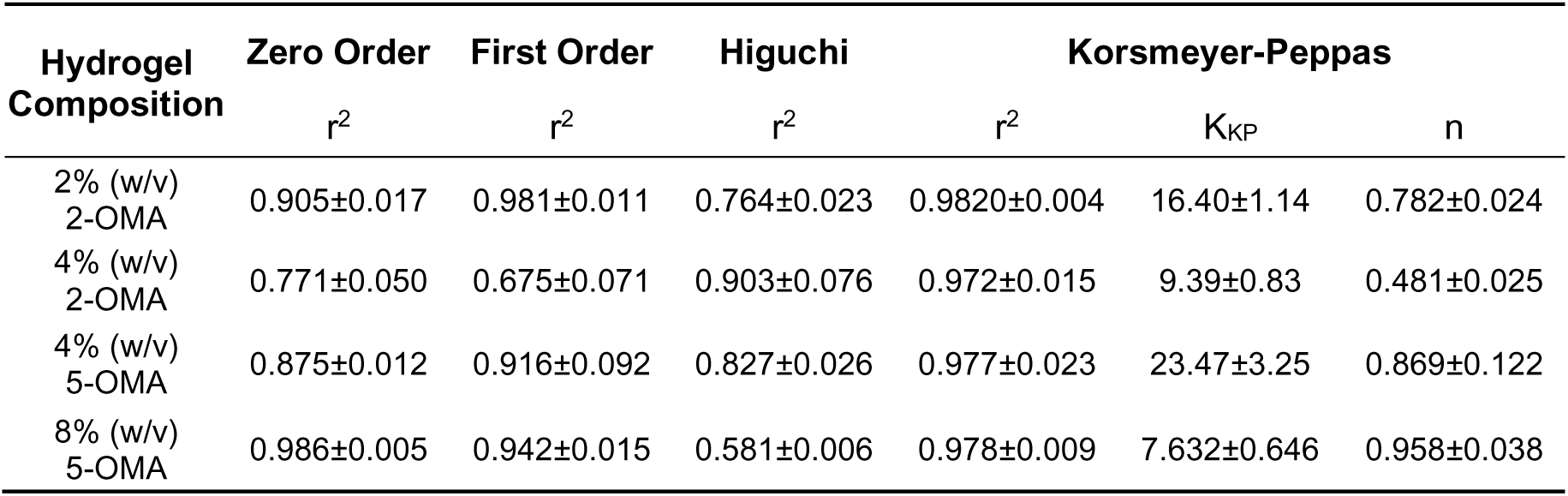
Regression coefficients obtained from the kinetic equations reflecting exosome release from OMA hydrogels.

| Hydrogel Composition | Zero Order | First Order | Higuchi | Korsmeyer-Peppas |  |  |
| --- | --- | --- | --- | --- | --- | --- |
| | $r^2$ | $r^2$ | $r^2$ | $r^2$ | $K_{KP}$ | $n$ |
| 2% (w/v) 2-OMA | 0.905±0.017 | 0.981±0.011 | 0.764±0.023 | 0.9820±0.004 | 16.40±1.14 | 0.782±0.024 |
| 4% (w/v) 2-OMA | 0.771±0.050 | 0.675±0.071 | 0.903±0.076 | 0.972±0.015 | 9.39±0.83 | 0.481±0.025 |
| 4% (w/v) 5-OMA | 0.875±0.012 | 0.916±0.092 | 0.827±0.026 | 0.977±0.023 | 23.47±3.25 | 0.869±0.122 |
| 8% (w/v) 5-OMA | 0.986±0.005 | 0.942±0.015 | 0.581±0.006 | 0.978±0.009 | 7.632±0.646 | 0.958±0.038 |

The diffusion/relaxation model (Korsmeyer-Peppas, KMP) exhibited high correlation coefficients (r^2^ > 0.97) for all compositions, suggesting it is the most suitable for studying exosome release from 2-OMA and 5-OMA hydrogels. The release constant (K_KP_) from the KMP model varied significantly across different compositions, with 4% (w/v) 2-OMA hydrogels showing the lowest value (9.39 ± 0.83) and 4% (w/v) 5-OMA hydrogels the highest (23.47 ± 3.25), reflecting differences in exosome release rates. Moreover, the K_Kp_ value showed a decrease with increasing macromer concentration in both types of hydrogels.

The release exponent values (*n*) obtained according to the KMP model for all compositions were above 0.45, ranging from 0.481 ± 0.025 for 4% (w/v) 2-OMA hydrogels to 0.958 + 0.038 for 8% (w/v) 5-OMA hydrogels, which suggests a non-Fickian diffusion mechanism. Accordingly, the release mechanism appeared to be a complex process that involves the degradation and swelling of the OMA hydrogels, as well as exosome diffusion.

### 3.3 Cytocompatibility and functionality of OMA hydrogels

The *in vitro* cytocompatibility of photocrosslinked 4% (w/v) 2-OMA and 5-OMA hydrogels was examined by fluorescence staining with a Live/Dead assay (**Fig. 4A**). As mentioned, HeLa cells were cultured using a Transwell culture plate where hydrogels were placed on the upper insert of the well and allowed to degrade over time. Since cells were placed in the lower chamber of the plate they were cultured in the presence of the polymer as it degraded into the culture medium.

**Figure 4.**
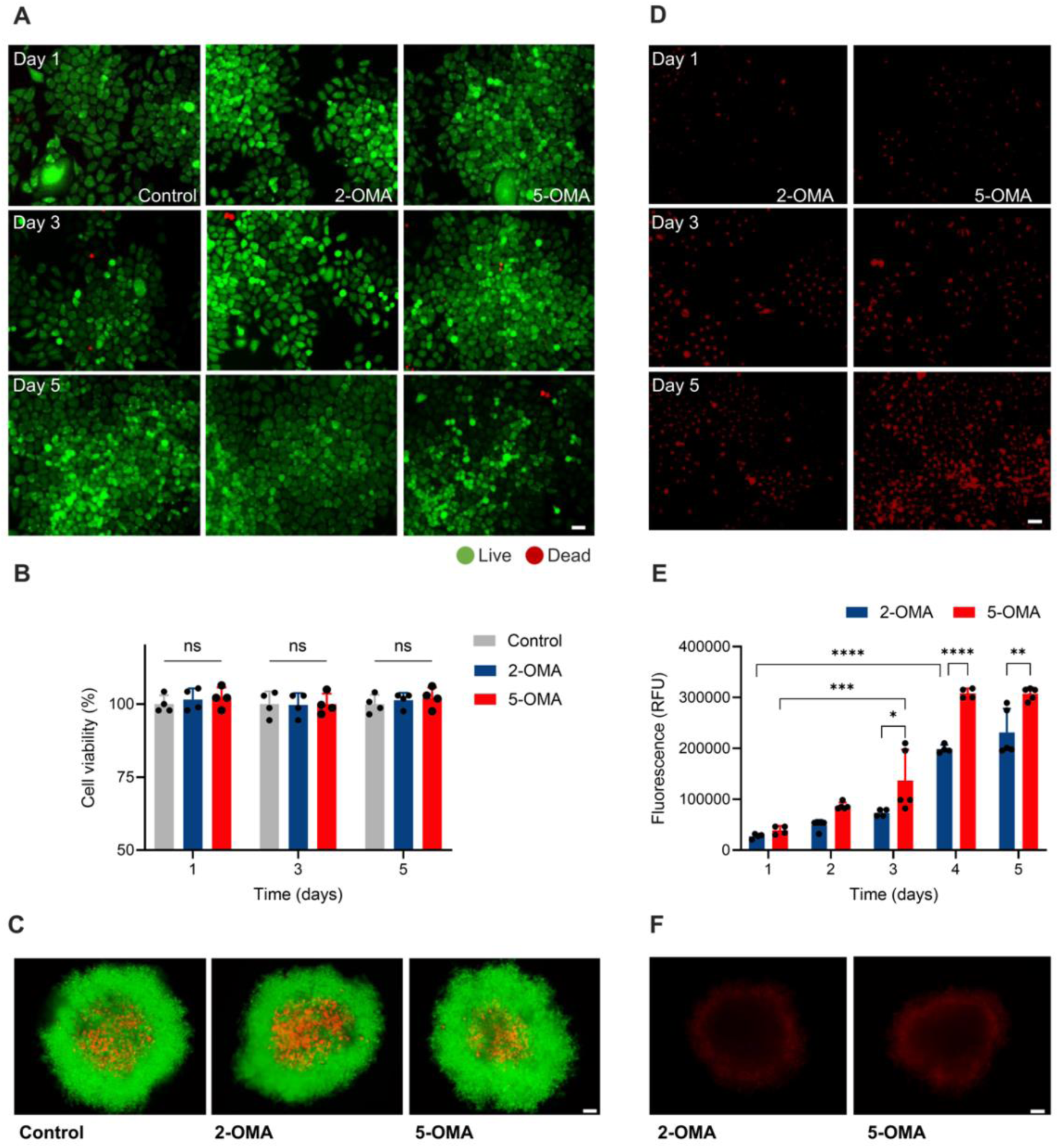
Biocompatibility and functionality of OMA hydrogels containing exosomes. **(A)** Live/Dead staining images of monolayer HeLa cells and **(B)** cell viability percentage of monolayer HeLa cells after culture with OMA hydrogels, normalized to untreated control cells (n=4). **(C)** Live/Dead staining images of HeLa spheroids encapsulated in OMA hydrogels after 3 days of culture. Images depicting exosome uptake by monolayer HeLa cells **(D)** cultured with OMA hydrogels encapsulating sEVs labeled with BODIPY-TR ceramide. **(E**) Quantification of exosome uptake as determined by the relative fluorescence intensity (RFU) of monolayer HeLa cells after culture with OMA hydrogels encapsulating fluorescently labeled exosomes. (F) Images depicting exosome uptake by HeLa spheroids after 3 days of culture within OMA hydrogels. Data are presented as the mean ± SD. Scale bar: 100 µm. Statistics analysis was performed using a two-way analysis of variance (ANOVA) with the Tukey significant difference post hoc test (\**p*<0.05; \*\**p*<0.005; \*\*\**p*<0.0005; ****p<0.0001; ns, not significant) in **(B)** and **(E)**.

The viability of HeLa cells cultured with the 2-OMA hydrogels was 96.87 ± 1.17%, 98.11 ± 1.01% and 97.35 ± 0.90% after 1, 3 and 5 days of culture, respectively. Likewise, the viability of cells cultured with 5-OMA was 97.11 ± 1.01%, 97.19 ± 1.56%, and 96.12 ± 2.15% after 1, 3, and 5 days of culture, respectively. These results indicate that the presence of either type of polymer does not significantly influence cell viability in a monolayer culture compared to controls grown without the hydrogels **(Fig. 4B)**.

In a separate set of experiments, HeLa spheroids were encapsulated within photocrosslinked 4% (w/v) 2-OMA and 5-OMA hydrogels and cultured over time. It was observed that, within these spheroids, some interior cells were dead while most peripheral cells were alive. This pattern of cell viability is expected during spheroid development, suggesting that factors such as UV light exposure, the crosslinking reaction and/or the degradation products of the OMA hydrogels do not negatively influence cell viability within this three-dimensional structure (**Fig. 4C)**.

To validate the delivery of exosomes from the OMA hydrogels, 4% (w/v) photocrosslinked 2-OMA and 5-OMA hydrogels containing fluorescently labeled exosomes were placed in the insert of a Transwell culture plate, while cells were cultured in the lower chamber. The fluorescently labeled exosomes were released from the OMA hydrogels and endocytosed by target cells (**Fig. 4D**).

Exosome uptake was quantified by measuring the fluorescence intensity of target cells grown in monolayer at predetermined time points (**Fig.4E**). Fluorescence intensity increased every day during culture, and it was sustained for 5 days. After 3 days, fluorescence intensity in cells cultured with the 5-OMA hydrogels was higher than in cells cultured with the 2-OMA hydrogels. Furthermore, fluorescence intensity in cells cultured with the 5-OMA hydrogels significantly increased after 3 days when compared to day 1, while a significant increase was observed after 4 days in cells cultured with the 2-OMA hydrogels.

In addition, HeLa spheroids were encapsulated within photocrosslinked 4% (w/v) 2-OMA and 5-OMA hydrogels containing fluorescently labeled exosomes and cultured for 3 days. Staining of spheroids within both 2-OMA and 5-OMA hydrogels indicated exosome uptake by HeLa cells (**Fig. 4F**). Therefore, exosome uptake is possibly related to the sEV release profile and/or the physicochemical properties of the OMA hydrogels. Altogether, these findings suggest that OMA hydrogels can release exosomes *in vitro*.

## 4. Discussion

Our results showed that oxidized and methacrylated alginate (OMA) is a suitable biomaterial to produce photocrosslinkable and biodegradable hydrogels for the release of exosomes. Alginate oxidation enhanced the degradability of the hydrogels, which may partly dictate the rate of exosome release. Accordingly, OMA hydrogels with a higher degree of oxidation, which undergo rapid degradation, showed rapid exosome release, while slowly degrading OMA hydrogels were highly stable and could retain exosomes for a prolonged period. This is significant for applications that require either rapid exosome release and clearance of the delivery vehicle or long-term exosome release by a sustained release depot [32].

As previously described, increasing the alginate oxidation degree results in enhanced hydrolysis of the OMA hydrogels [47]. The degradation properties of these hydrogels correlated with their swelling and mechanical properties, which were also tunable by varying macromer concentration. Here, the rapid degradation of the 5-OMA hydrogels increased exosome release, while the more slowly degrading 2-OMA hydrogels showed enhanced exosome retention.

The degradation and release rates observed in OMA hydrogels show an important relationship between their physicochemical properties and functionality. Specifically, the chemical composition, crosslinking density, and the resulting internal network structure of the gels, are crucial in determining their swelling ratio, and directly influence their rheological properties [48].

Indeed, the storage modulus (G’) and loss modulus (G”) result from the internal structure of the gels and the environmental conditions, which are also determining factors of their swelling behavior [49,50]. Understanding the interplay between these properties is crucial for controlled release applications, where the release rate of bioactive factors may be controlled by diffusion through the internal polymer network [51]. Our findings also agree with previous studies that show a correlation between the storage and loss moduli of gels and bioactive factor release rate [52]. These studies have shown that hydrogel stiffness may regulate the exosome release behavior and could play an important role in the clinical applications of these biomaterials [37,53].

Here, the difference in the alginate oxidation degrees for OMA hydrogels affects the availability of degradation sites [29]. While both hydrogels underwent photocrosslinking because of their methacrylation, a lower alginate oxidation degree results in a network that is less prone to hydrolytic degradation due to fewer oxidized groups. This contributes to the formation of a comparatively stronger network in terms of mechanical strength and stability, as observed in the increased stiffness and slower degradation rates of the 2-OMA hydrogels when compared to those with a higher degree of oxidation. Similarly, increasing the macromer concentration could facilitate the formation of a more densely crosslinked hydrogel network.

Recent reports have presented the development of unmodified and chemically modified alginate, as well as alginate-based for the controlled delivery of EVs [25,26,54]. These studies show that exosomes can be retained for prolonged periods in alginate-based hydrogels, making them suitable for their sustained delivery [55–57]. However, controlled release of these vesicles is not expected from alginate hydrogels, because the release of bioactive molecules is generally thought to be controlled by diffusion, a process driven by concentration gradients between the hydrogel and its surrounding environment [58].

Accordingly, in the absence of specific interactions between the hydrogel and the encapsulated exosomes, or stimuli-responsive release triggers, the diffusion process may not allow for modulation of the release rate in response to specific delivery requirements. Furthermore, alginate is a highly hydrophilic polymer, which results in a high-water content, facilitating the rapid diffusion of encapsulated molecules out of the hydrogel [59,60]. This makes alginate hydrogels prime systems for nutrient transport in tissue engineering applications. However, it poses a challenge for applications requiring controlled release.

With this issue, Huang *et al.*, modified alginate by the addition of an RGD motif to restrain the EVs within the formed hydrogels [26]. This strategy aims to introduce specific interactions between the hydrogel and encapsulated exosomes, overcoming the proposed release mechanism driven by diffusion. In this work, we observed controlled release of exosomes from OMA hydrogels, suggesting that the chemical modifications introduced to the alginate may significantly influence the release mechanism beyond simple diffusion.

Alginate oxidation and methacrylation may influence the network structure, degradation behavior, and interactions with encapsulated exosomes, contributing to a more controlled release [61]. Furthermore, these chemical modifications result in photocrosslinkable hydrogels with tailored degradation rates. Controlled degradation can lead to a more predictable and sustained release of exosomes, as the OMA hydrogels degrade at a designed rate, gradually releasing the encapsulated exosomes. These modifications also influence the physicochemical properties of the hydrogels, such as crosslinking density and mechanical strength. These factors determine the diffusion path for the encapsulated exosomes, directly affecting the release kinetics [62].

In this context, kinetic modeling of the exosome release data showed that it followed the Korsmeyer and Peppas model, showing that release kinetics exponentially corelate with fractional release over time [63]. The release exponent indicated a non-Fickian transport mechanism in which hydrogel composition influences the exosome release profile [64]. This also suggests that exosome release from OMA hydrogels may not only be dictated by diffusion but that it also involves the degradation and swelling of the hydrogel.

Specifically, swelling of the hydrogel alters its porosity and structure, affecting the diffusion pathway and rates of exosome release [65]. This relationship indicates that exosome release from OMA hydrogels may be driven by both the physical changes of the gels and the following modulation of diffusion dynamics. Consequently, the release process may be greatly influenced by the hydrogel structure, which can be readily tuned by varying the degree of alginate oxidation and macromer concentration.

An optimal delivery system would release exosomes at a predetermined rate for a specified period. Thus, tuning the physicochemical properties of OMA hydrogels provides a starting point to control the release rate. In this context, more complex hydrogel systems that incorporate additional functionalities or components, such as stimuli-responsive elements, multiple materials and crosslinking capabilities, or other nanomaterials may provide a more precise control over the release mechanisms [66]. However, whether these more complex designs offer superior control compared to more simple strategies, warrants further comparative studies.

Indeed, the aim to reduce the complexity of the biomaterial while still controlling the release rate is highlighted by the need for practicality, cost-effectiveness, and ease of manufacturing for specific applications [67]. For instance, if rapid scaling or regulatory compliance is critical, then simple systems that balance functionality with manufacturability become highly valuable.

It is worth mentioning that the *in vitro* release assays performed here were conducted under highly controlled conditions and the release data may not correlate to the *in vivo* performance of this systems [68]. Furthermore, this work only examined the performance of OMA hydrogels at physiological pH and no mechanical forces were applied to the system during degradation. Similarly, the *in vitro* degradation and swelling assays may also vary from the *in vivo* performance of hydrogels. However, the potential applicability and tunability of OMA hydrogels has been previously demonstrated for *in vivo* applications [47]. Furthermore, photocrosslinking allows rapid and convenient polymerization that has previously shown no effect on bioactive factor integrity or cell viability [36].

Our findings show that neither the photocrosslinking process nor the products of OMA degradation influenced the viability of target cells, as determined through the use of a viability stain. This results underscore the high cytocompatibility (i.e., not eliciting toxic effects on cells) of this biomaterial, as demonstrated by previous research [29,31]. To further support the applicability of OMA hydrogels in a clinical context, comprehensive assessments including cell proliferation and metabolic activity could be proposed. Such assessments, focusing on particular cellular processes may be needed to fully understand the interaction of OMA hydrogels with cells in targeted therapeutic contexts. Notably, these evaluations have been conducted in previous studies, suggesting its suitability for therapeutic delivery and its potential for broad clinical applications [29,31,69].

This work extended into the functional dynamics of these systems, demonstrating that exosomes released from OMA hydrogels can be endocytosed by target cells. The exosomes uptake assays performed in this study primarily reflect the endocytosis of these vesicles. Although this may not be considered functional because exosomes are not exerting a specific functional effect, for the purpose of this study, the uptake of these EVs by target cells is sufficient for establishing the groundwork for future research projects. These studies may be required to better understand the full spectrum of exosome functionality for different clinical purposes.

Previous studies on the use of EVs as therapeutics have explored a wide range of delivery mechanisms, such as oral administration, intravenous infusion, inhalation, topical application, and intraperitoneal injection [70–72]. However, these studies show non-specific accumulation of EVs in a single tissue, alongside rapid uptake by macrophages and off-target effects. Localized and controlled delivery systems, such as those based on OMA, may improve the pharmacokinetics of exosomes and selectively deliver them to target tissues to treat a wide range of medical conditions.

A notable approach within this context is the development of injectable hydrogels for the targeted delivery of cancer therapeutics. These hydrogel systems aim to suppress tumor growth by the local delivery of therapeutics that may cause serious side effects if administer systemically [73–75]. For example, Chao *et al*., developed an alginate-based delivery system for localized chemotherapy with immune adjuvants that can be combined with conventional intravenous injection of immune check point inhibitors [76]. These strategies have shown great potential for clinical translation and support the exploration of including exosomes into these systems for drug delivery and antitumor immunity [77–79]. Hydrogel-based local delivery of immune checkpoint blockade therapy, immune cell therapy, tumor vaccines, and oncolytic viruses are emerging as promising modalities of anticancer therapy that also facilitates combination therapies [80–82].

In this context, OMA has shown to be injectable, potentially enabling direct injection into a target site and facilitating rapid gelation through photocrosslinking, which may be performed *in situ* [24,36]. OMA with a higher oxidation degree can be proposed as a rapidly degrading hydrogel for the rapid release of therapeutic exosomes to prevent cancer recurrence by hydrogel implantation at the surgical site after resection, to avoid malignant transformation of non-malignant cells, to hinder tissue invasion at a metastatic site and to boost the local immune response [77,83]. On the other hand, slowly degrading OMA can be proposed for applications that require prolonged release, such as long-term local delivery of modified exosomes that may replace numerous chemotherapy cycles by sustaining a consistent therapeutic concentration in the target site. Moreover, the continuous exposure to therapeutic agents, efficiently delivered by these EVs, could inhibit the development of resistant mechanisms, offering a potential strategy to prevent therapy resistance in multiple types of cancer [81,84,85].

Further research is required to produce OMA hydrogels with a defined release mechanism for optimal exosome delivery for particular applications. A major limitation to these proposed applications is that OMA is not adherent to tissues. However, formulations combining OMA with poly(ethylene glycol)(OMA/PEG) have shown enhanced and tunable adhesion strengths [31]. These bioadhesive properties may not only facilitate surgical procedures, such as tissue grafting or wound closure, but also suggest that OMA hydrogels could achieve direct tissue adherence, potentially avoiding the need of a secondary dressing material or extensive chemical modifications before *in vivo* implantation.

Finally, although there is some understanding of the immediate biological compatibility of hydrogels systems, comprehensive studies of their long-term effects after implantation, such as immunogenic responses and the potential of chronic inflammation or fibrosis are needed [86,87]. These studies should aim not only to assess the potential adverse effects arising from these processes but also to explore strategies to reduce or prevent them.

## 5. Conclusion

In this work, we developed tunable, photocrosslinkable and biodegradable OMA hydrogels for exosome delivery. The swelling, degradability and rheology of OMA hydrogels were tunable by adjusting the alginate oxidation degree and macromer concentration. By controlling the degradation behavior of the hydrogels, accelerated exosome release was achieved increasing the alginate oxidation level. Conversely, a lower alginate oxidation degree enhanced exosome retention and long-term stability of the hydrogels. OMA hydrogels showed remarkable cytocompatibility and were able to deliver exosomes to target cells *in vitro*. The strategy of tuning the physicochemical properties of OMA hydrogels to control exosome release might be applied to several biomedical applications, such as targeted drug delivery, regenerative medicine, and tissue engineering. Research of value beyond the scope of this work include further refinement of release mechanisms for EV delivery, adhesion to biologic tissue and long-term *in vivo* safety evaluations. Likewise, multiple clinical trials are currently evaluating the toxicity and the ability to halt or reverse disease progression of exosome-based therapeutics. This study highlights the possibility of developing hydrogel systems for the localized, controlled release of exosomes. Exploring the combination of various delivery strategies represents a promising direction for future research. While this study focused on tuning the physicochemical properties of OMA hydrogels for exosome release, future studies could integrate these findings with additional strategies, such as targeted cell delivery or stimuli-responsive release mechanisms to improve the efficacy of therapeutic applications.

Progress on the development of these systems may shed light on their specific applications by demonstrating their versatility and efficacy in localized and targeted therapeutic strategies. The ability to control the release kinetics of exosomes from hydrogels means that treatments can be tailored to the specific needs of different diseases and patients. Furthermore, the fine-tuning of the properties of OMA hydrogels can be leveraged to gain a better understanding on how these systems interact with different tissues and disease models, leading to the identification of conditions where these systems offer the most significant therapeutic advantage.

Similarly, comprehensive *in vivo* studies may reveal the long-term safety of these systems, which is key for applications involving implantable devices or long-term treatment regimens. Demonstrating how OMA hydrogels can be combined with other treatment strategies, such as conventional drug therapy, gene therapy, or immunotherapy could result in the development of novel multidisciplinary approaches. Indeed, progress in these systems may not only highlight their diverse clinical applications but also their potential for improving patient care.

## 6. Declarations

## Acknowledgments

The authors would like to thank the UIC-Tec de Monterrey seed funding program for the initial funding in the development of this work. Seed funding was provided by the Richard and Loan Hill Department of Biomedical Engineering at University of Illinois Chicago and by the School of Engineering and Science at Tecnológico de Monterrey. Authors at Tecnológico de Monterrey thank the School of Engineering and Science and the FEMSA-Biotechnology Center for their support through the Molecular and Systems Bioengineering Research Group. Some figures were generated using BioRender.com.

## Authors Contributions

**Sergio Ayala-Mar**: Conceptualization (supporting); Formal Analysis; Investigation; Writing – Original Draft Preparation. **Oju Jeon**: Conceptualization (supporting), Investigation; Methodology; Supervision (supporting), Writing – Review and Editing. **Pedro Chacón-Ponce**: Formal Analysis; Investigation. **Eben Alsberg**: Conceptualization (lead); Funding Acquisition; Methodology; Project Administration; Supervision (lead); Writing – Review and Editing. **José González-Valdez**: Conceptualization (lead); Funding Acquisition; Methodology; Project Administration; Supervision (lead); Writing – Review and Editing.

## Data availability statement

Derived data supporting the findings of this study are available from the corresponding authors upon reasonable request.

## Ethical approval statement

Not applicable.

## 8. Supplementary Information

**Figure S1.**
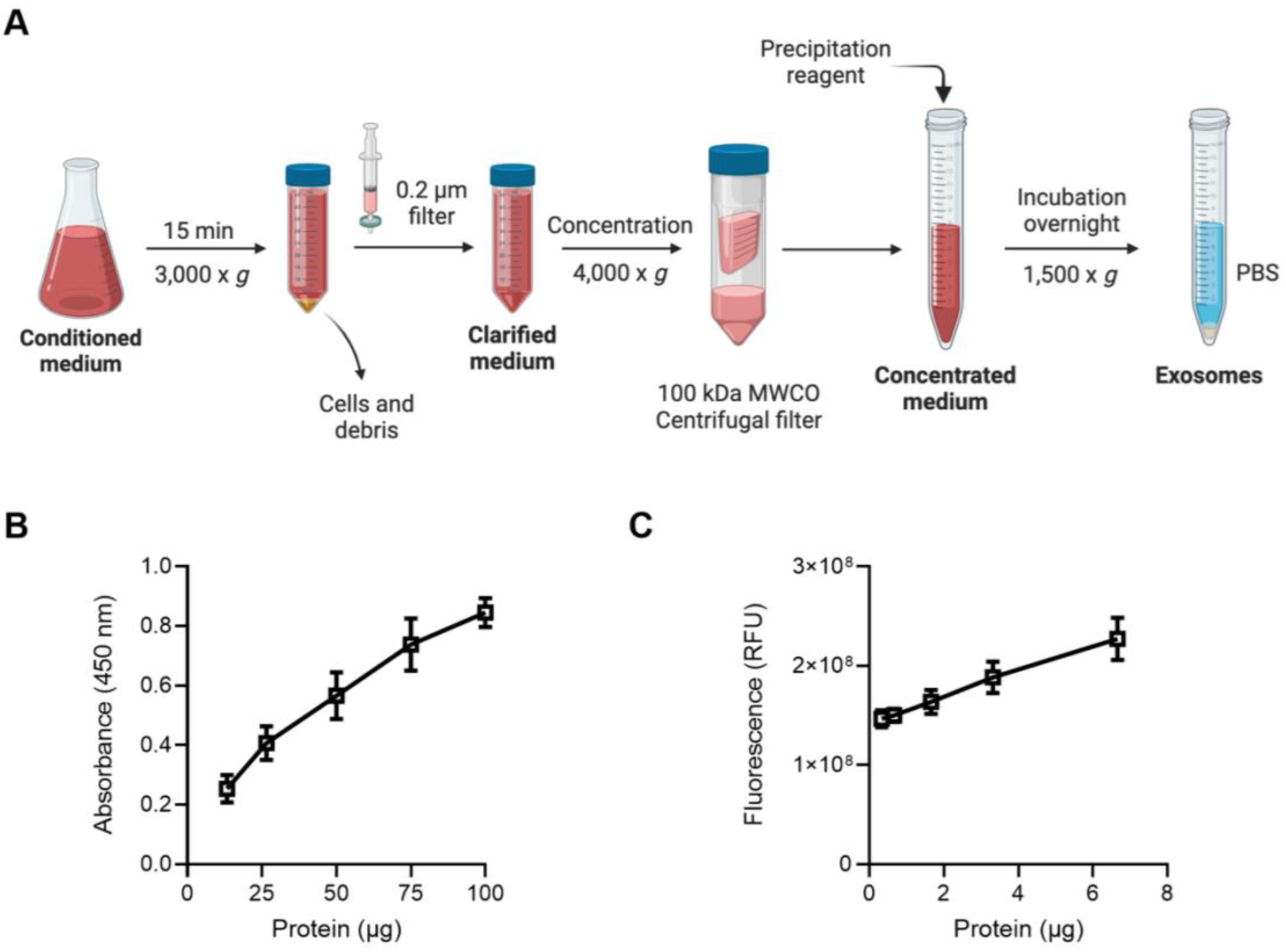
Exosome isolation process and validation by ELISA for CD63 detection and acetylcholinesterase (AChE) activity assay. **(A)** Schematic representation of the exosome isolation protocol from conditioned cell culture medium, beginning with centrifugation at 2,000 x *g* to remove cells and debris, followed by 0.2 µm filtration. The clarified medium is then concentrated using a 100 kDa molecular weight cutoff (MWCO) centrifugal filter and mixed with a precipitation reagent. Following overnight incubation, the mixture is centrifuged at low speed to collect an exosome pellet. **(B)** ELISA results for CD63 in the exosome stock demonstrate a positive correlation between absorbance at 450 nm and protein content, confirming the presence of the tetraspanin CD63. **(C)** AChE activity assay results illustrate an increase in relative fluorescence units (RFU) with increasing protein content, indicative of exosomal enzymatic activity. Data are presented as mean ± standard deviation (n=3).

**Figure S2.**
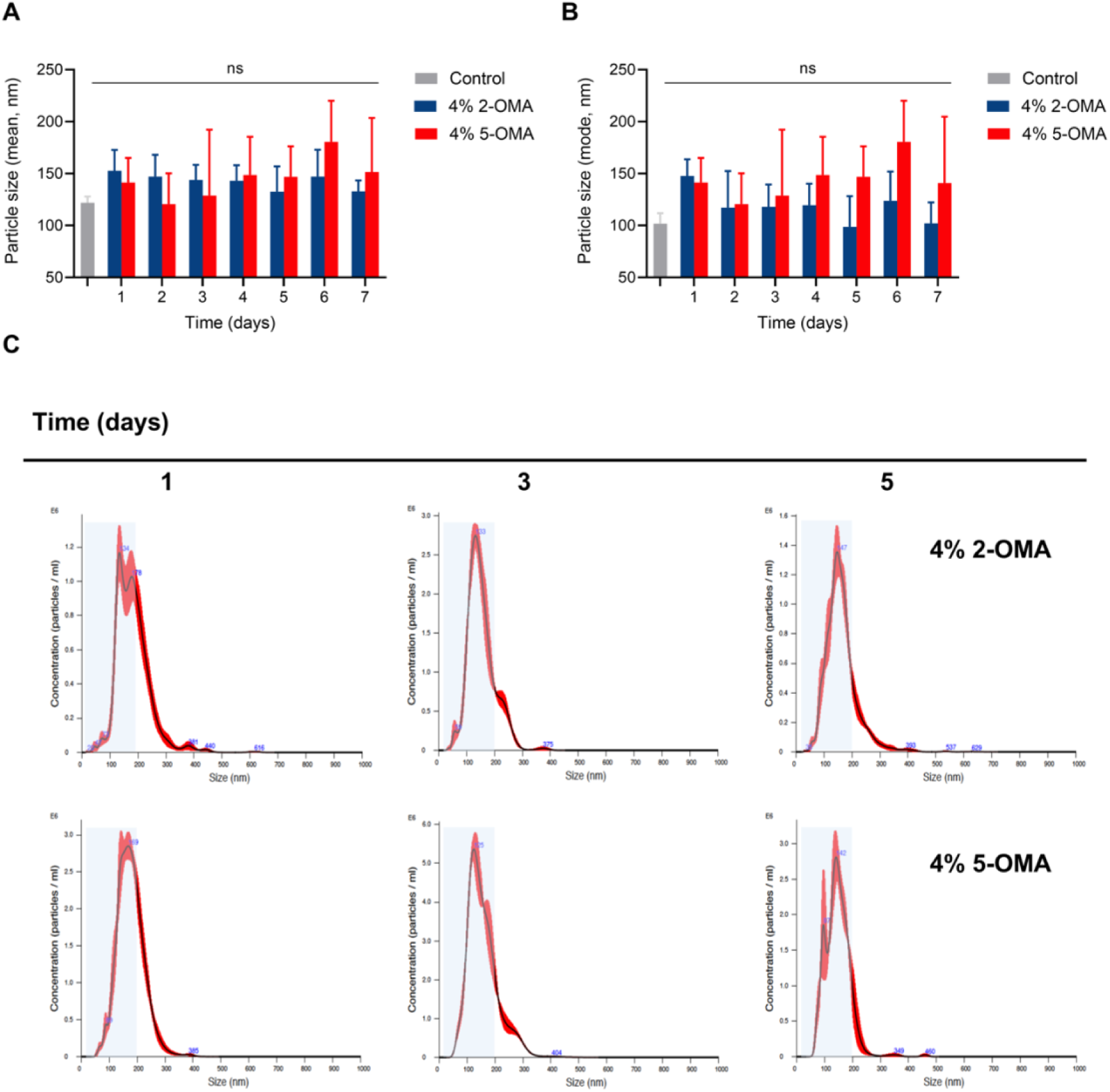
Analysis of exosome particle size and distribution after being released from 2-OMA and 5-OMA hydrogels over time. **(A)** Mean particle size of exosomes incorporated into 2-OMA and 5-OMA hydrogels, measured over a 7-day period using nanoparticle tracking analysis (NTA). Exosome release from OMA hydrogels is compared against the control (exosome stock). **(B)** Mode particle size of exosomes released from the 2-OMA and 5-OMA hydrogels, as well as the control, over the same period. **(C)** Representative size distribution profiles of exosome samples recovered from the hydrogels on days 1, 3 and 5. Shaded regions in blue indicate a particle size range of 10-200 nm. Statistical significance in A and B was assessed using multiple unpaired t-tests with a False Discovery Rate (Q) of 1% for comparisons between all groups. Data are presented as the mean ± standard deviation (n=5); ns indicates not statistically significant.

